# Seasonal changes in coat colour and sexual size dimorphism in a subtropical ungulate

**DOI:** 10.1101/2025.02.10.637565

**Authors:** Tania A. Perroux, Alan G. McElligott, George M. W. Hodgson, Kate J. Flay

## Abstract

Phenotypes reflect how organisms adapt to their environments. Hong Kong (HK) feral cattle, a crossbreed of *Bos taurus taurus* and *Bos taurus indicus*, present an opportunity to study these adaptations in one of the very few global cattle populations not directly controlled by humans. These cattle are free-ranging since their release from farms in the 1970’s. HK has a subtropical climate, characterized by high humidity and temperatures during the wet season, and scarce precipitation during the dry season. We studied seasonal coat colour changes in HK feral cattle, and sexual dimorphism in body size and horn length. We provide the first evidence of seasonal changes in coat colour in cattle, with paler coats being more common in the wet season, while darker coats prevailed in the dry season. These seasonal changes were influenced by temperature, humidity, solar radiation, and body condition. We found that males were larger and had longer horns than females. Our results show a male-biased sex dimorphism in the HK feral cattle. Additionally, our findings suggest that thermoregulation costs drive colouration in these cattle. The phenotypic plasticity we demonstrate in these subtropical feral cattle improves our knowledge of the adaptations of ungulates to their habitat.

## INTRODUCTION

Animal phenotypes are visible characteristics of individuals that result from the interplay of genotype and environment. In wild populations, in which genetic and ecological information may be scarce, phenotypes provide an overview of the adaptations of organisms to their environment. Phenotypic traits may differ from previous populations as a result of human-generated selection pressures (Stockwell *et al*., 2003; Allendorf & Hard, 2009) and changes in climatic conditions (Jones *et al*., 2018; Oli *et al*., 2023; Otte *et al*., 2024). A number of phenotype characterization methods have been developed for livestock and companion species (*e.g.* Amin, 2009; Hameed et al., 2022; Ouchene-Khelifi et al., 2018), where individuals can be easily restrained for measurements. However, in populations where restraint is more challenging (*e.g*. large ungulates, wild populations), phenotyping data is often assessed using corpses (Fulgione *et al*., 2016; Schulz *et al*., 2024) or using invasive procedures (*e.g.* capture; Canale et al., 2016). Non-invasive methodologies, such as measures on photographs (*i.e.* photogrammetry), are becoming more popular for phenotypic data collection in free-ranging animals (*e.g.* Bro-Jørgensen & Dabelsteen, 2008; Cambreling et al., 2025; Morandi et al., 2022; Prinsloo et al., 2021), providing valuable information about understudied populations.

Coats serve multiple functions, including providing thermal insulation by creating a barrier between the skin and the ambient environment, as well as reflecting solar radiation to reduce heat gain (Stuart-Fox *et al*., 2017; Beltran *et al*., 2018; Zimova *et al*., 2018). Generally, paler coats reflect more solar radiation compared to darker ones, which is advantageous for animals in tropical environments (Stuart-Fox *et al*., 2017). Conversely, darker coats provide protection against ultraviolet radiation (UV; Newell et al., 2021). As postulated by Gloger’s rule (reviewed by Delhey, 2019), species inhabiting hot and wet environments, such as tropical and subtropical habitats, are expected to have darker coats (Cloudsley-Thompson, 1999; Wacker *et al*., 2016). This trend has been confirmed in various mammals, notably primates (Kamilar & Bradley, 2011), wild pigs (Newell *et al*., 2021), rodents (Ge *et al*., 2021) and more generally in artiodactyls (Stoner, 2003) by comparing populations from temperate habitats to populations in tropical and subtropical habitats.

Phenotypic traits can exhibit plasticity, as demonstrated by the periodic moulting of coats in various endotherm species (Beltran *et al*., 2018; Zimova *et al*., 2018). Moulting is influenced by external factors, with photoperiodism (*i.e.* daylight duration) triggering the initiation of the process, while climate conditions, such as temperature and snow cover, affect the speed of moulting (Zimova *et al*., 2018). Additionally, the timing and duration of moulting are governed by internal factors, including age, sex, body condition, and reproductive status, all modulated by the endocrine system (Beltran *et al*., 2018; Zimova *et al*., 2018). Rapid adaptations in coat colouration are often observed in species expanding their distribution (*e.g.* Newell et al., 2021). Predation pressure is a primary driver of coat colouration, as camouflage decreases predation risk through background matching (Atmeh et al., 2018; Caro, 2005). In environments with significant seasonal contrasts, such as temperate and arctic habitats, these seasonal changes in colouration facilitate adaptations that improve concealment in response to fluctuating vegetation and soil conditions (Zimova *et al*., 2014, 2018). Typically, endotherm coats are darker in summer and lighter in winter, with variations in colour reflecting the balance of eumelanin (black/brown pigments) and phaeomelanin (yellow/red pigments). Seasonal changes in coat colour are expected when the energetic costs of producing a new coat are lower than the costs associated with suboptimal colouration (Beltran *et al*., 2018).

Sexual size dimorphism typically results from intrasexual competition and/or intersexual mate choice (Lukas & Clutton-Brock, 2014; Clutton-Brock, 2017; Shuker & Kvarnemo, 2021) and can be expressed in a variety of measurements of body lengths and secondary sexual traits (*e.g.* horns). Although recent meta-analysis revealed that sexual size dimorphism is not as common as previously reported (Tombak *et al*., 2024), artiodactyl (order Artiodactyla) males tend to be both larger and longer than females (McPherson & Chenoweth, 2012; Cassini, 2020; Tombak *et al*., 2024). In polygynous ungulates, males are larger than females due to intrasexual competition, as larger body size and longer horns in males result in higher social status and fitness (McElligott *et al*., 2001; Knierim *et al*., 2015; Cassini, 2020). Several bovid species are characterized by striking sexual dimorphism, particularly in body size (Cassini, 2020) and horn shape (Capellini & Gosling, 2006.; Caro et al., 2003; Nasoori, 2020), while others have little to no sexual dimorphism (D’Ammando *et al*., 2022). More extreme sexual dimorphism decreases longevity in male bovids (Bro-Jørgensen, 2012). Horns are secondary sexual characteristics displayed by either one or both sexes in bovids (Davis *et al*., 2014). Most bovids grow horns continuously by the proliferation of germinal layer cells (Davis *et al*., 2014). The horns have a diversity of important biological functions, including defence from predators, fighting weapons, courtship and thermoregulation (Emlen, 2008; Stankowich & Caro, 2009; Davis *et al*., 2014).

There are 1.5 billion cattle globally, with the vast majority managed on farms (FAO, 2021). It is almost impossible to study cattle phenotypes in populations that are not directly controlled by humans, as there are only a few feral/wild populations. Although it is known that darker cattle display higher signs of heat stress and at lower thresholds than paler coated cattle in agricultural settings (Santos *et al*., 2017), evidence in wild/feral populations is lacking, and seasonal coat colour changes have not been previously documented in cattle. The feral cattle of Hong Kong (HK) are a genetically distinct mix of *Bos taurus taurus* and *Bos taurus indicus*, with some evidence that they also carry genes from wild Asian bovids (Barbato *et al*., 2020), similar to other Southeast Asian cattle (Zhang *et al*., 2007; Gao *et al*., 2017; Xia *et al*., 2023; Luo *et al*., 2025). Originally used as draught animals, these cattle are now regarded as part of the local heritage (SKBW, 2017). HK feral cattle are likely to have adapted to match the local environment and were subjected to low selection pressure for production traits even when they were used on farms (Barbato *et al*., 2020). They have striking phenotypic diversity, although their appearance has not been formally studied (Barbato *et al*., 2020).

Despite the striking phenotypic diversity of these animals, little is systematically known about the HK feral cattle phenotypes. We aimed to assess their phenotypes in a non-invasive way, with a particular focus on body size, horn length and coat colour in respect to sexual dimorphism and seasonality. We expected the HK feral cattle to follow the general assumption of sexual dimorphism in artiodactyls, with males being larger and having longer horns than females. Additionally, we predicted distinct seasonal changes in coat colour between the dry and wet seasons, influenced by solar radiation as observed in other species. In the absence of major predation pressure on this population, we hypothesized that observed seasonal coat colour changes would be driven by thermoregulation needs and that changes in body condition may reflect the costs of moulting.

## METHODS

### 1. Seasonal Coat Colour

#### 1.1 Study sites and animals

For the 317 cattle scored for sexual size dimorphism, we also recorded coat colour. We then selected 12 herds for longitudinal scoring over a period of 13 months, with a total of 253 cattle scored longitudinally. These herds were selected based on their relative ease of access, ensuring reliable follow-up, while also maintaining good representation of the population (*i.e.* herd size, sex ratio, environment type, provisioning, proximity and frequency of interactions with humans). Following initial scoring (*i.e.* August 2022 during body size measurements), we visited each of the 12 selected herds once a month from January 2023 to January 2024 to obtain individual coat colour scores over time. A total of 2,371 coat colour scores were recorded, with each individual scored 1 to 14 times (mean = 9.37 ±SD 4.43).

#### 1.2 Measures

We scored coat colour while maintaining a distance of at least 10 meters to the cattle. Coat colour was scored as a qualitative and subjective measure, controlling for lighting conditions and observer subjectivity influences by using distinct categories of score (Bro-Jørgensen & chart consisted of five coat colours: pale, red, grey, dark and black (Fig. 1). All coat colours were estimated by a single observer to reduce subjectivity of scoring, similar to Agosta et al. (2017). To ensure reliability of scores, scores made in August 2022 were also checked by three independent observers, and scores were discussed until all observers reached an agreement. This allowed us to refine the coat colour chart (Fig. 1). We used this colour chart to assess three body parts: fore, middle and hind (D’Ammando et al., 2022; Loehr et al., 2008). Then, we used the mode of those three body part coat colours to obtain each individual’s coat colour per month. Individuals for which coat colour in all three body parts differed were not retained for that month; this was the case for eight individual cattle scores. Further description of the eight data points removed is presented in Table S1. Moulting is known to follow patterns across the body which are well described in several species (Zimova *et al*., 2018), but such prior knowledge was not available for our study animals. Hence, these discrepancies of colour across body part may simply reflect the moulting process but could not be analysed in the present study. To assess coat colour changes, coat colours were replaced by an ordinal scale, with palest coats scored 1 and darkest scored 5 (Fig. 1).

**Figure 1:**
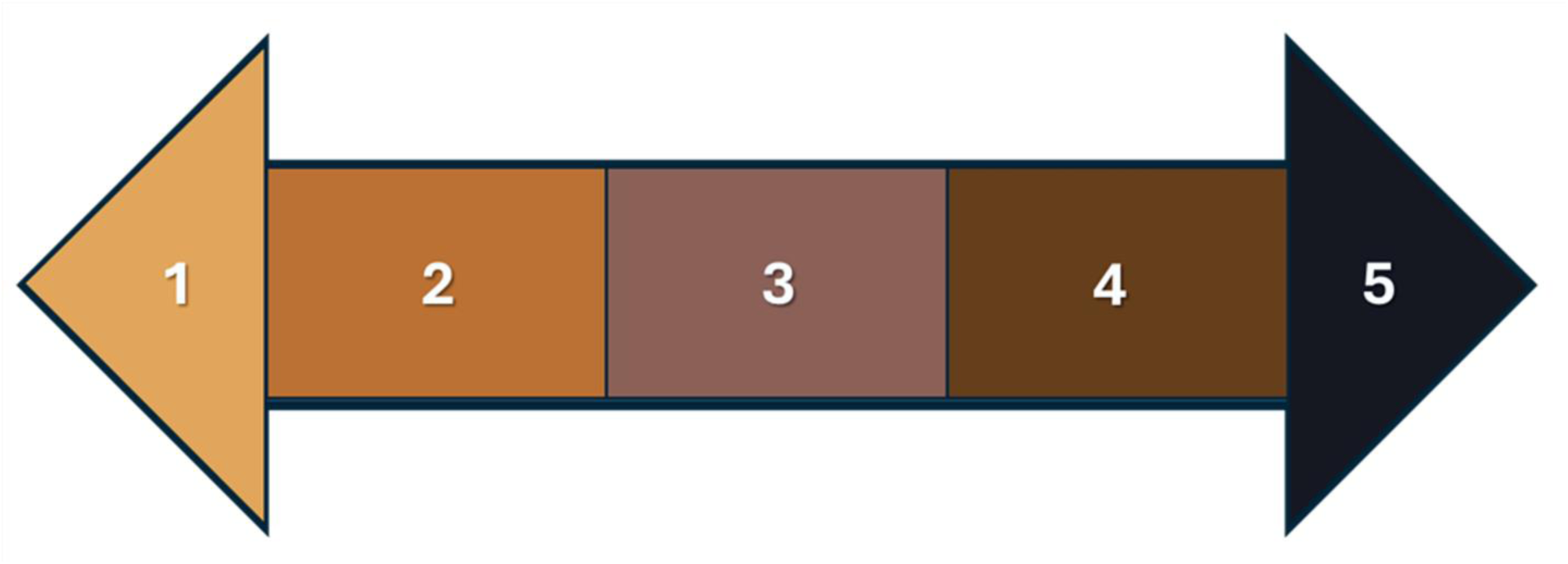
Coat colour scale used to characterize coat colours in Hong Kong feral cattle with 1 corresponding to palest coats, 2 for coats scored as red, 3 for grey, 4 for dark and 5 black (the darkest coat colour). This colour scale is adapted from Stoner (2003).

Each individual received a Body Condition Score (BCS) at the time of coat colour scoring based on a modified visual 9-point scale, with 1 being the thinnest individuals (low body condition), and 9 the fattest (high body condition). This scale was adapted to the HK feral cattle based on beef cattle BCS (D’Occhio *et al*., 2019; Weik *et al*., 2021). To control for the subjective nature of body condition scoring, training was conducted to ensure good scorer reliability (for six weeks from April 1st 2022 to May 5th 2022), with body condition scored once a week by a single observer on cattle in a single herd (n = 60 individuals). BCS scores were expected to be highly correlated from one week to the next, and therefore the scorer trained until BCS from one week to the subsequent reached a significant correlation of more than ρ = 0.8 (based on Spearman’s rank correlation) for three consecutive weeks. This same observer scored all BCS used in this study.

#### 1.3 Climate Data

We extracted monthly averages for mean daily temperature (in C°), relative humidity (%RH), total rainfall (in mm), total bright sunshine duration (in hours) and mean daily global solar radiation (in megajoule per square meter) from the Hong Kong Observatory database (HKO, 2020). Seasons were defined based on a Principal Component Analysis of climate data from HKO from January 2015 to July 2022, hence independent from our data collection period. Using a Hierarchical Cluster Analysis, we confirmed that HK subtropical weather followed three seasons: dry season (from December to February); the wet season (from May to October); and the intermediate season (March, April and November). Climate during our sampling period is summarized in Table S2.

#### 1.4 Statistical analysis

As our data derived from normality (as tested by Shapiro-Wilk test), we analysed the repeatability of scores between the years with a paired (per individual) Mann-Whitney signed-rank test, comparing scores made in August 2022 with scores in August 2023, and scores made in January 2023 with scores in January 2024. Differences in distribution of coat colours between the sexes were analysed using a χ² test and adding months as a layer in the contingency table. Coat colour was transformed into a continuous scale from −4 to 4 to form the variable “coat colour with negative values for coats getting paler and positive values for coats getting darker while 0 indicates no changes in coat colour.

The distribution of coat colours in the population was analysed using a General Linear Model for Ordinal Logistic Regression (OLR) using the function “clm” (package “ordinal”). A first OLR compared the distribution of coat colours to the magnitude of coat colour change. Then, a second OLR was used to test the impact of climate variables and BCS from the month of the score and the month before the score. A backwards elimination procedure was conducted, with variables with the highest p-values removed stepwise until only significant variables remained. Multicollinearity of the predictors was checked using the Variation Inflation Factor (VIF) with the function “vif” (package “car”). A Brant test was conducted to ensure that the proportional odds assumption was respected, using the “brant” function (package “brant”), with p-values above 0.05 indicating that the assumption holds. Predicted Probabilities (PP) were extracted from the OLR model to facilitate interpretation of the results, using meaningful points on significant predictors (*i.e.* using five integers along the observed predictor natural range).

Coat colour change was analysed using an analysis of covariance (hereafter ANCOVA) with the R function “aov” (package “stats), from which significance was obtained using the function “Anova” specifying for type III (package “car”). Coat colour change was used as the dependent variable (outcome). Sex and season were entered as independent variables in the ANCOVA. Climate variables from the month of the measurements and the previous month were added as covariates and eliminated following a backwards elimination procedure. Specifically, covariates with the highest p-values were removed stepwise until only significant covariates remained. ANCOVA assumptions were checked as follows. Homoscedasticity was checked using a Bartlett test, with p-values above 0.05 indicating that the assumption is respected. The homogeneity of regression slopes was obtained by modelling every possible interaction effect between a fixed factor and a covariate. The assumption was respected if no interaction effect had a p-value above 0.05. Due to our large sample size, we did not have to conform to normality assumptions based on the central limit theorem (Islam, 2018). Specifically, with a sample of 253 cattle, we can assume a precision of 5-7% (Islam, 2018). Post-hoc tests were conducted for significant independent variables using a Bonferroni correction.

### 2. Sexual Size Dimorphism

#### 2.1 Study sites and animals

The HK feral cattle population is currently estimated at 900 individuals (Pinkham *et al*., 2022) found in a variety of environments, from urbanized areas to rural country parks, with some herds on geographically isolated islands. Their main predators, the South China tiger (*Panthera tigris tigris*) and leopard (*Panthera pardus*), have not been observed in HK since the 1940’s, and although cases of predation on calves by Burmese python (*Python bivittatus*) and feral dogs have

Conservation Department (AFCD) of the Government of the Hong Kong Special Administrative Region (HKSAR) provides veterinary care to injured individuals, but routine care is not provided (AFCD, 2018). The AFCD aims to achieve a stable cattle population primarily relying on reproduction control via surgical sterilization (Pinkham *et al*., 2022).

We studied 30 different cattle herds (Fig. 2), scoring one to 43 individuals in each herd (Table S3), with a total of 317 individuals scored (about a third of the population). We visited each location at least once and up to three times. Herds were distributed over 11 different country parks (n=5 to 117 individuals per country park), with country parks defined by the AFCD. We collected data during Hong Kong’s wet season, between 27^th^ of July 2022 and 30^th^ of October 2022.

**Figure 2:**
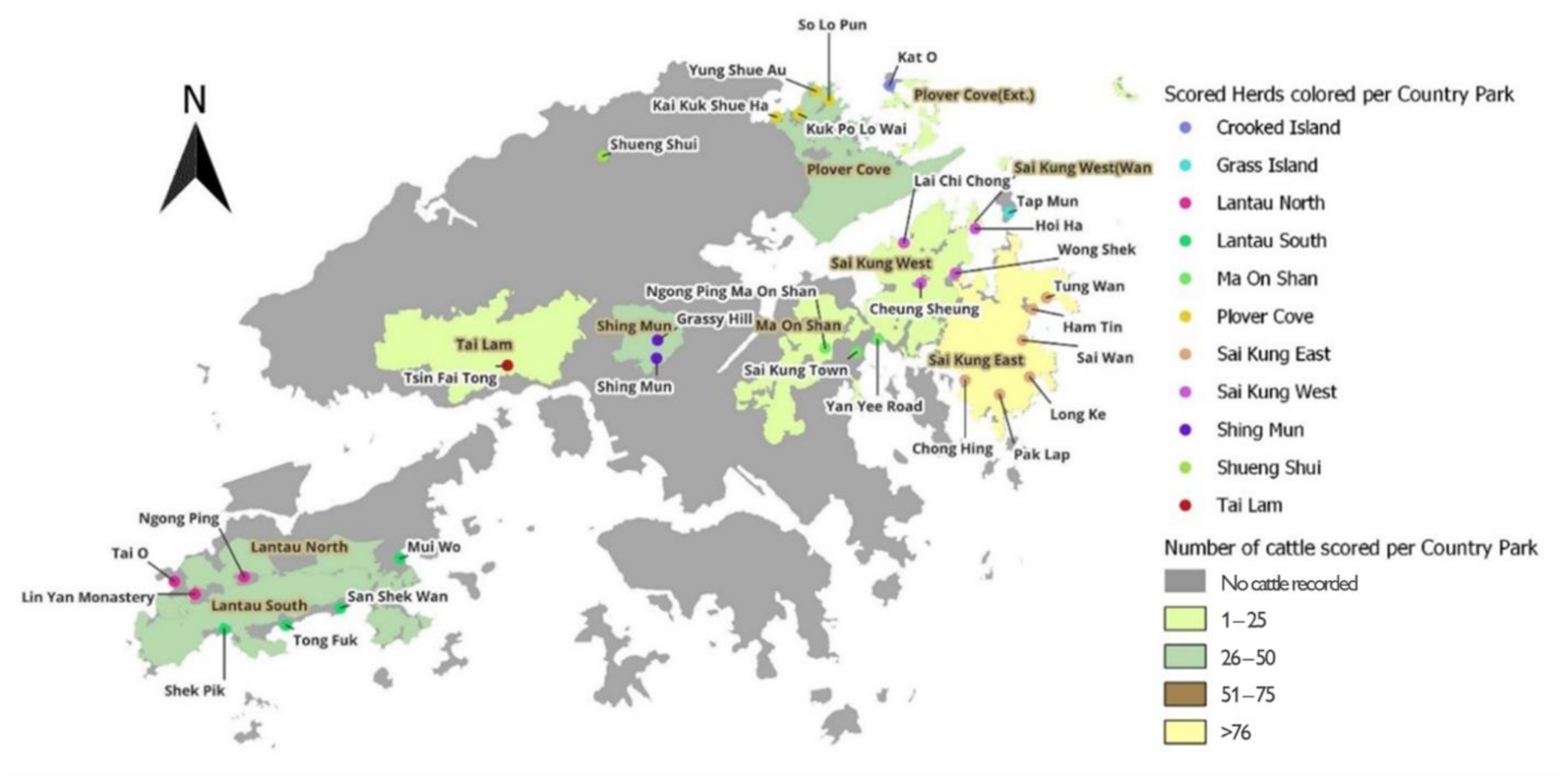
Map of the herds sampled for body size measurements in Hong Kong SAR. The colours represent the number of cattle scored in country parks. Country parks were defined following the Agriculture and Fisheries Conservation Department (AFCD) of Hong Kong SAR. Country Park names have brown labels, while white labels indicate herd names. Dot colour indicates country park membership. Map was created using the QGIS software (version 3.28.2).

We excluded calves and only scored adult cattle as recommended for description of breed populations (FAO, 2011). Our exclusion criteria defined calves following two criteria: *i.* absence of apparent horns or horns detached to the skull (horn tips moving separately to jaw movements) and *ii.* similar appearance to same-aged individuals (Fig. S1). We scored all cattle meeting the inclusion criteria (*i.e.*, adult cattle) and gave each a unique identifier; either its tag as provided by the AFCD, or the name of the herd followed by the number of individuals found that day (*e.g.* the 4^th^ individual found in Tai O was identified as TO4), unless the animal had been named for other studies.

#### 2.2 Measures

We scored horn length by visual comparison to ear length along three categories: shorter, similar or longer than ear length. We developed this method to enable scoring horn phenotypes without restraining individuals. We accounted for broken and asymmetrical horns by adding them as a category, resulting in a total of five categories. Horn length was scored for all study animals (170 females and 147 males). Side view photos were taken of each individual to assess body size measures (Ferreira & Funston, 2010; Berger, 2012; Tarugara *et al*., 2019).These photos could be taken without restraint, and digital analysis of photographs also decreases intra and inter-observer variability compared to measures made with a tape, particularly for linear measures (Goodenough *et al*., 2012). We used five measures: wither height, hip height, body length, hip width and chest depth (Tasdemir et al., 2011; Zhang et al., 2018; Table 1; Fig. 3). To ensure reliability of the measures, we used a triangulation method: the scorer took a photo of the cow and a field assistant. The distance to the animal was decreased slowly until equal distance between cow, scorer and field assistant was met, at which point the scorer took the photo. As soon as the photo was obtained, the scorer stepped away from the cow. As this was not possible for all individuals (i.e., not able to safely approach and/or the animal repeatedly moved away), body size measures were not recorded for all cattle, with measurements obtained for 122 females and 100 males. Using the photos, we took measures with Prime Ruler Android application (Grymala, 2022, version 1.7.6). We used the real height of the assistant divided by the height given by the app as a factor to correct the application measures as follows:

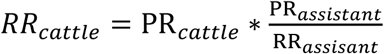

*where RR = real measure estimation and PR = Prime Ruler measurements*

**Figure 3:**
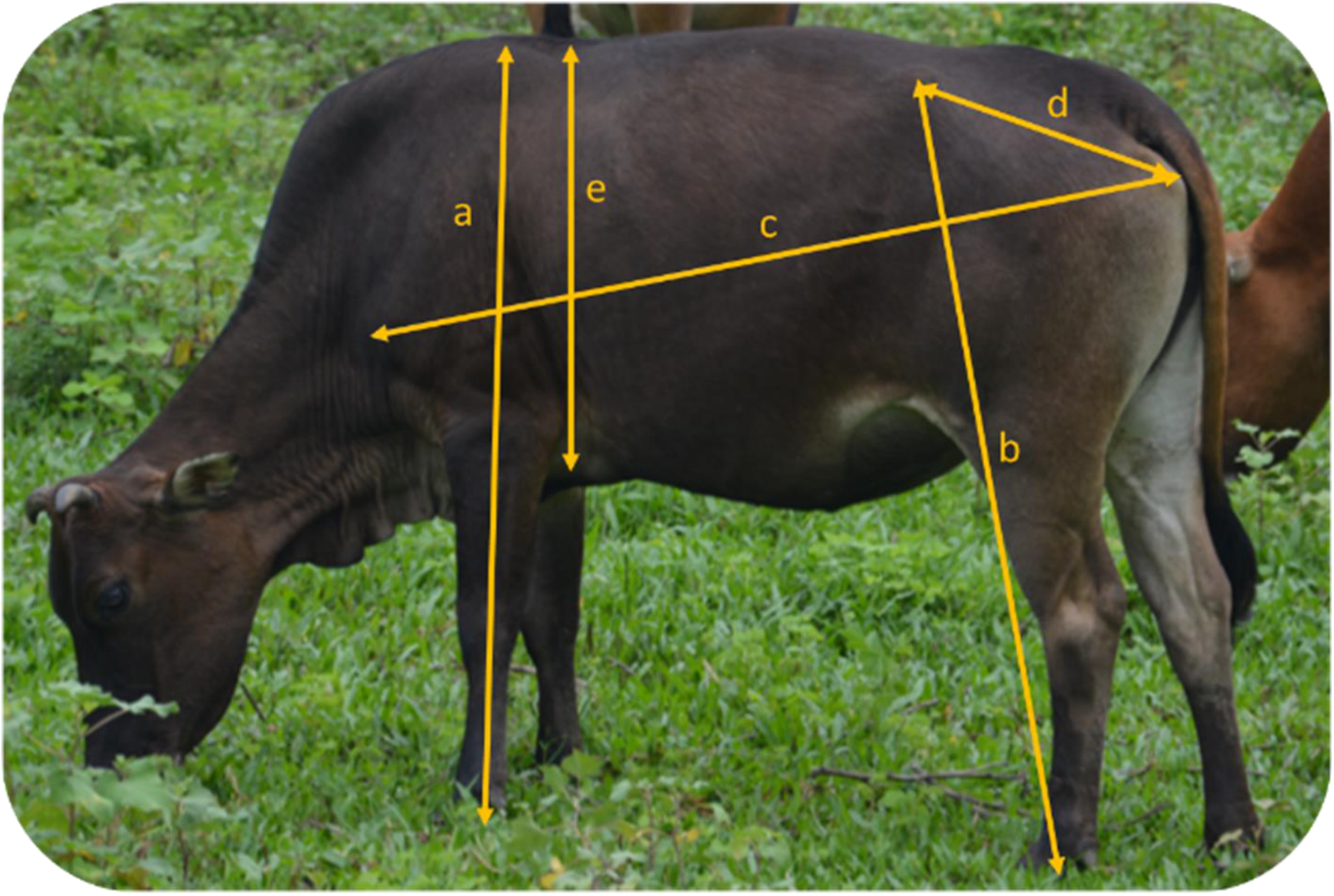
Different measures used to assess body size based on pictures of Hong Kong feral cattle: Wither Height (a), Hip height (b), Body length (c), Hip width (d) and Chest depth (e).

**Table 1:** Body size measures used to describe the Hong Kong feral cattle.

| Body Size Measure | Anatomical definition |
| --- | --- |
| Wither Height | From wither to hoof of fore limb |
| Hip height | From tuber coxae to hoof of hind limb |
| Body length | From shoulder joint to tuber ischii |
| Hip width | From tuber coxae to tuber ischii |
| Chest depth | From withers to brachial pit of fore limb |

#### 2.3 Statistical Analysis

We performed descriptive statistics using JASP software (JASP Team, 2022, Version 0.19.0.0), while further tests were performed using R (R Core Team, 2020, version 4.3.3). All scripts and datasets are provided on OSF database.

As all measures of body size were highly correlated (based on Spearman’s correlation, with all correlations ρ>0.54 and p<0.001, Table S4), we used a Principal Component Analysis (PCA; function “PCA”, package “FactoMineR”, Lê et al., 2008) to obtain a summary numerical variable for body size. The first Principal Component (PC1) of the PCA was selected, as it explained 77.81% of the variance, had an eigenvalue of 3.89, and was significantly positively correlated to all five measures of body size (p<0.0001). We used a Linear Mixed-effect Model (LMM; function “lmer”, package “lme4”, Bates et al., 2015) to assess for sexual dimorphism of body size; body size was used as the dependent variable and sex as the independent variable. Location nested within country park was used as a random factor. The significance was obtained using Satterthwaite’s method for single term deletions (“drop1” function, package “stats”).

Sexual dimorphism index (SDI) is a measure developed by Lovich and Gibbons (1992) to quantify the body size difference between the sexes. It was adapted for ruminants (Polák & Frynta, 2009, 2010; Cassini, 2020) as follows: 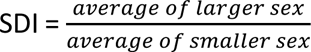. In our analysis, male was the larger sex used in the formula. This measure was calculated to describe sexual dimorphism in all five body measurements. Sexual dimorphism of horn length was assessed using a Pearson’s Chi-squared test (function “chisq.test”, package “stats) to compare the distribution of each category of the variable horn length between males and females.

### 3. Ethical Statement

Our research was reviewed and approved by the Animal Research Ethics Sub-Committee of the authors’ university (Internal Reference: A-0826). We undertook fieldwork in compliance with the Guidelines for the Treatment of Animals in Behavioural Research and Teaching (ASAB/ABS, 2022).

## RESULTS

### 1. Seasonal Coat Colour

Red coats are the most common coat colour in all seasons, while grey coats are the least common (χ² = 594.28, p <0.0001; Table 2; Fig. 4). Overall, 31.52% (95%CI = 29.33 – 33.80) of individuals changed coat colour throughout the year, though extreme changes are uncommon. While our methodology would identify individuals that go from black to pale (or pale to black), this was never observed, and extreme changes (−3 and 3) account for less than 1% of individual changes. Coat colours follow seasonal patterns, with no difference between the scores made in August 2022 and 2023 (W = 553.50, p = 0.40) and in January 2023 and 2024 (W = 471.00, p = 0.98), indicating that fluctuation of coat colour follows yearly cycles.

**Figure 4:**
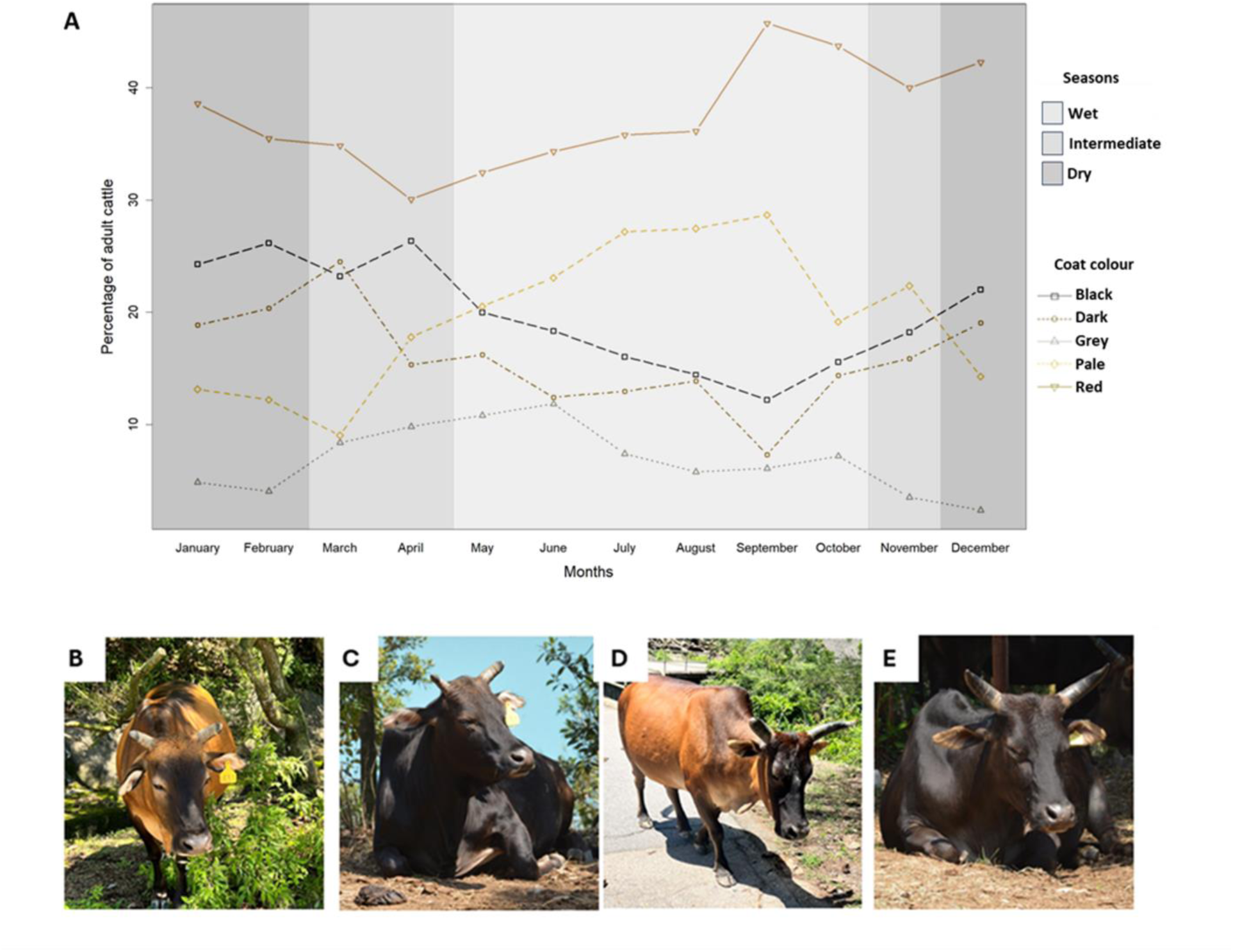
Monthly variation of coat colour of the Hong Kong cattle (A) and examples of coat changes within a same individual: female cattle (tagged 431) in the wet (B) and in the dry (C); and male cattle (tagged 17) in the wet (D) and in the dry (E). Coat colour was scored based on a colour chart.

**Table 2:** Coat Colour distribution in the Hong Kong feral cattle in the dry, intermediate and wet season, scored based on a colour chart.

| Season | Coat Colour | Counts | Frequency (in %) | 95%CI |
| --- | --- | --- | --- | --- |
| Dry<br>(686 scores) | Red | 267 | 38.92 | 35.34 - 42.62 |
|  | Black | 167 | 24.34 | 21.28 - 27.69 |
|  | Dark | 133 | 19.38 | 16.60 - 22.51 |
|  | Pale | 91 | 13.26 | 10.93 - 16.00 |
|  | Grey | 28 | 4.08 | 2.83 - 5.83 |
| Intermediate<br>(487 scores) | Red | 171 | 35.11 | 31.00 - 39.45 |
|  | Black | 110 | 22.58 | 19.09 - 26.50 |
|  | Dark | 90 | 18.48 | 15.28 - 22.16 |
|  | Pale | 81 | 16.63 | 13.58 - 20.19 |
|  | Grey | 35 | 7.18 | 5.21 - 9.83 |
| Wet<br>(1184 scores) | Red | 449 | 37.92 | 35.20 - 40.72 |
|  | Black | 190 | 16.04 | 14.06 - 18.24 |
|  | Dark | 156 | 13.17 | 11.36 - 15.22 |
|  | Pale | 295 | 24.91 | 22.53 - 27.45 |
|  | Grey | 94 | 7.93 | 6.53 - 9.61 |

There are significant differences between male and female coat colour distribution (χ² = 156.20, p <0.0001). Females are in general paler, with red being the most common coat colour in females (44.25%, 95%CI = 41.59 - 46.95), while black coats are the most common in males (30.80%, 95%CI = 28.07 - 33.67). These differences are not consistent throughout the year, with significant sexual differences in coat colour distribution between males and females in all months (p < 0.05, Table S5), except in May (χ² = 8.63, p = 0.07), June (χ² = 2.96, p = 0.56), September (χ² = 6.46, p = 0.16) and October (χ² = 8.92, p = 0.06).

Individuals with darker (black and dark) coat colours are more likely to change coat colour than cattle with red, grey and pale coats (OLR; OR = 1.80, 95%CI = 1.57 - 2.06, p <0.0001). Coat colour is predicted by the monthly average of the mean daily temperature from the month before the measurement (OLR; OR = −0.06, 95%CI = −0.08 - −0.05, p <0.0001, Table S6) and by the BCS at the time of the measurement (OLR; OR = −0.21, 95%CI = −0.33 - −0.10, p = 0.0001, Table S7). Following months of lower temperature (15 °C), the likeliness for cattle to be darker increases (PP; dark = 0.19; black = 0.30), while following higher temperatures (35 °C), the likeliness of cattle being paler increases (PP; pale = 0.32; red = 0.40). Similarly, the likeliness of having a paler coat increases in individuals with higher body condition (BCS 7, PP; pale = 0.28; red = 0.39), while in individuals with lower body condition (BCS 3) the likeliness to be darker increases (PP; dark = 0.17; black = 0.23). The interaction between BCS and the temperature from the previous month could not be computed as it increased VIF/multi-collinearity above 60, making it unsuitable for interpretation.

The likeliness for coat colour to change is significantly explained by seasonal variations (F_(5,1647)_ = 5.71, p = 0.003), with coat colours getting paler in the wet season while they get darker in the dry season (Fig. 4). Specifically, cattle are more likely to change colour between the intermediate and the wet season (Tukey Post-hoc, t = −3.14, p = 0.005, Table S8). There is no difference between males and females in the magnitude of changes in coat colours (F_(5, 1647)_ = 0.37, p = 0.54). The impact of seasonal variation in coat colours is driven by the average humidity (F_(5, 1647)_ = 17.44, p<0.001) and the mean daily global solar radiation (F_(5,1647)_ = 33.61, p<0.001) in the month of scoring.

### 2. Sexual Size Dimorphism

Males are significantly larger than females (LMM, p = 0.002, Fig. 5) in all body size measurements as indicated by the SDI (Table 3). While male and female cattle are dimorphic, their dimorphism is moderate, with a mean SDI of 1.07, ranging from 1.05 for hip height and body length to 1.1 for chest depth.

**Figure 5:**
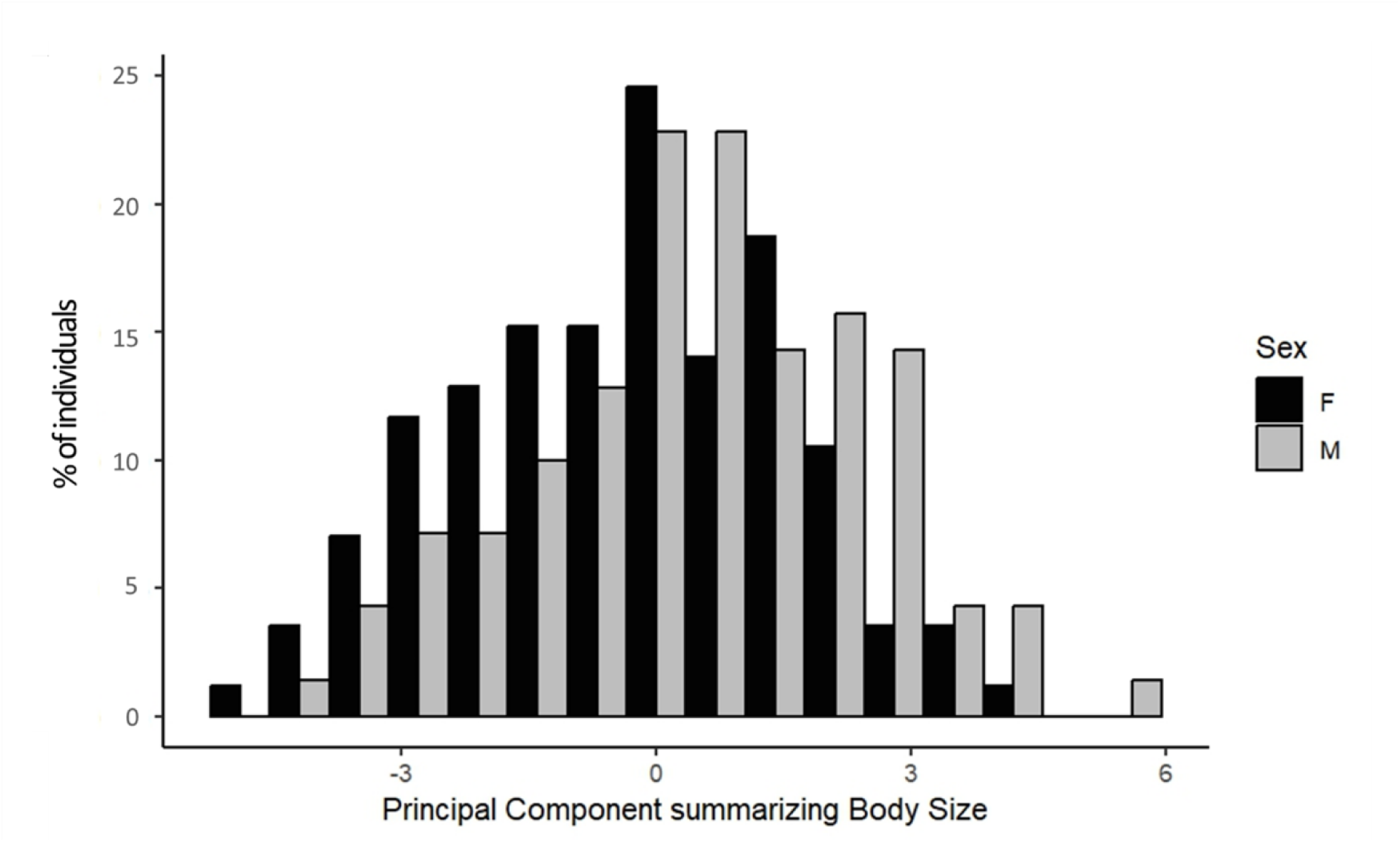
Body size of 100 male (light grey) and 122 female (black) Hong Kong feral cattle. Body size is expressed as a Principal Component (PC, unitless) obtained from a Principal Component Analysis (PCA) summarizing five measures of body size: wither height, hip height, body length, hip width and chest depth.

**Table 3:** Body size measurements (expressed in centimetres) and Principal Component (PC, unitless) obtained from a Principal Component Analysis (PCA) summarizing the five measures of body size for 100 male and 122 female Hong Kong feral cattle. Sexual Dimorphism Index (SDI) is given for all five body size measurements as the difference between the sexes (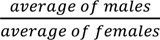

|  |  | Mean | Minimum | Maximum | SDI |
| --- | --- | --- | --- | --- | --- |
| Wither Height | F | 101.65 | 66.34 | 146.01 | 1.08 |
|  | M | 110.58 | 67.96 | 156.21 |  |
| Hip height | F | 95.20 | 51.29 | 144.50 | 1.05 |
|  | M | 100.88 | 30.92 | 139.08 |  |
| Body length | F | 112.10 | 70.42 | 161.19 | 1.05 |
|  | M | 118.48 | 49.19 | 174.87 |  |
| Hip width | F | 30.55 | 12.67 | 61.22 | 1.07 |
|  | M | 32.97 | 10.16 | 52.39 |  |
| Chest depth | F | 55.54 | 7.14 | 91.35 | 1.1 |
|  | M | 61.43 | 14.51 | 92.40 |  |
| PC1 | F | -0.33 | -5.56 | 4.75 |  |
|  | M | 0.38 | -4.77 | 6.10 |  |

**Figure 6:**
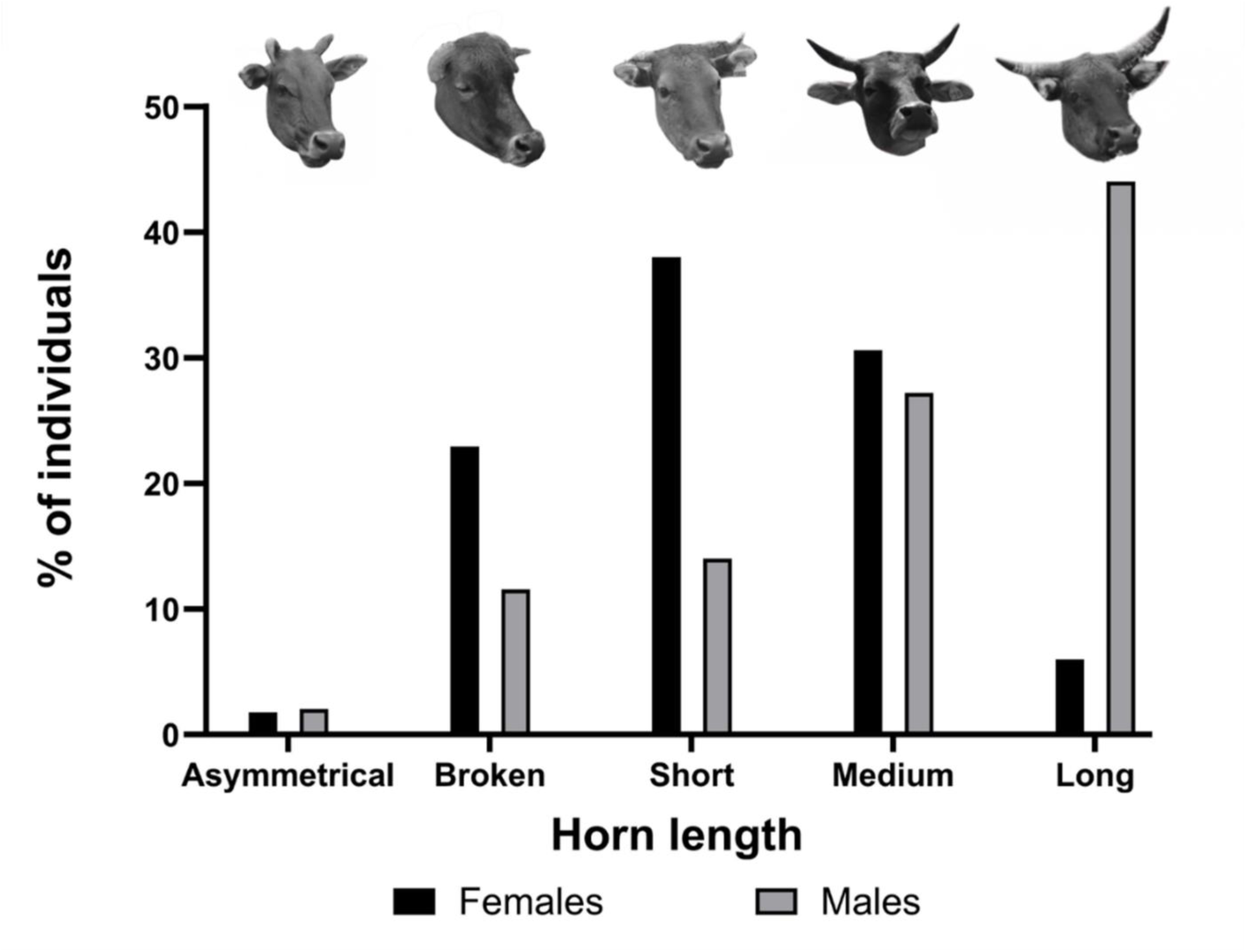
Distribution of horn lengths in 147 males (grey) and 170 females (black) Hong Kong feral cattle. Horn length was measured visually by comparing it to ear length.

Males have longer horns than females (χ²=68.52, p <0.0001; Fig. 6). While short horns are the most common horn type in females (38.24%, 95%CI = 21.26 - 45.72), long horns are the most frequent horn type in males (44.22%, 95%CI = 36.43 - 52.29). Asymmetrical horns are uncommon in both males (2.04%, 95%CI = 0.69 - 5.82) and females (1.76%, 95%CI = 0.60 - 5.05). Females are more likely than males to have broken horns (22.94%, 95%CI = 17.26 – 29.81, and 11.56%, 95%CI = 7.34 – 17.73, respectively).

## DISCUSSION

We provide the first phenotypic analysis of sexual dimorphism in body size and horn length in the Hong Kong (HK) feral cattle and quantify their seasonal coat colour changes. HK feral cattle are sexually dimorphic, with males being darker, larger and having longer horns than females. Additionally, females are more likely to have broken horns than males. We provide the first evidence of seasonal coat colour variation in cattle, with a third of individuals showing coat colour changes every month. Coat colours were predicted by body condition and the temperature from the previous month, while the magnitude of seasonal coat colour change was predicted by humidity and solar radiation. As global temperatures rise and weather patterns shift, many species are predicted to face important imbalances in thermoregulation (*e.g.* North et al., 2023), driving phenotypic expression to adapt to increasingly unpredictable environmental conditions. By understanding the pressures prey populations undergo in the absence of predators and the key role of phenotypic plasticity in their ability to adapt to changing environment, we can improve population dynamics projections.

HK cattle exhibit seasonal changes in coat colour, with more individuals displaying paler coats in the wet season and darker coats in dry season. Thermoregulation appears to be the primary driver of seasonal coat colour changes in HK cattle, influenced by humidity and solar radiation, as seen in other prey species facing high energetic costs from climate factors (Zimova *et al*., 2014; Kennah *et al*., 2023). Darker coats are generally better suited to tropical and subtropical environments, aligning with Gloger’s rule (Kamilar & Bradley, 2011; Delhey, 2019; Ge *et al*., 2021). However, the thermal melanism hypothesis predicts that lighter coats can help regulate heat gain and water loss in hot and humid climates (Delhey, 2019; Marcondes *et al*., 2020). For instance, in springbok *(Antidorcas marsupialis),* individuals with paler coats thrive in summer, while those with darker coats are better suited to winter climates (Hetem *et al*., 2009). Heat stress is a significant welfare concern in farmed cattle, negatively impacting behaviour and physiology, which can compromise long-term health when climatic conditions exceed cattle thermal neutral range (Cook *et al*., 2007; Becker *et al*., 2020; North *et al*., 2023). Darker cattle breeds tend to suffer more from excessive heat loads compared to breeds with paler coats (Mader *et al*., 2006; Brown-Brandl *et al*., 2006). Similarly, coat colour variation within breeds leads to individual differences in heat load, with black coated Holstein cattle having higher surface temperature than white cattle (Façanha *et al*., 2010; Santos *et al*., 2017). Our findings show that darker individuals have lower body condition, suggesting a higher cost associated with darker coats, in alignment with previous findings in dairy cattle (Anzures-Olvera *et al*., 2019). Humidity is expected to significantly impact thermoregulation costs in tropical populations (Marcondes *et al*., 2020), which supports our observation that humidity drives the magnitude of seasonal changes in coat colour. Genes related to UV protection and heat tolerance have been identified in other Chinese cattle breeds (Lei *et al*., 2006; Gao *et al*., 2017; Xia *et al*., 2023). Overall, our results support the thermal melanism hypothesis, as we found that temperature, humidity, and solar radiation influence seasonal coat colour changes.

Our findings provide the first evidence of seasonal changes in coat colour in a subtropical cattle population. While coat colour changes in cattle have been previously associated with age (Schmutz, 2012) and with nutritional deficiencies, with for instance copper deficiency being associated with paler coats in farmed cattle (Olkowski & Wojnarowicz, 2018). The association we found between darker coats and lower body condition highlights that it is unlikely to be related to nutritional deficiency and likely associated with seasonal patterns instead. Additionally, repeatability of scores across the years supports the idea of seasonal patterns of coat colour changes in HK cattle. In tropical ungulates, paler coats could be expected in the dry season for camouflage against vegetation, while darker coats in the wet season provide background matching for darker soil and foliage (see for instance Newell et al., 2021). However, our findings do not support these expectations. Coat colour variability is influenced by a trade-off between thermoregulation and predation risk (Beltran *et al*., 2018; Zimova *et al*., 2018; Kennah *et al*., 2023). As HK feral cattle were farmed (and thus likely protected by farmers) when predators were still present in HK, there may have been little to no pressure from predation risk on this population historically. Hence, coat colouration of HK cattle may not serve a function as an anti-predatory response, resulting in it not matching the background colours as would be expected in other free-ranging ruminants with high predation pressure.

We found that darker coats were present in all seasons, although these were more common in males and in the dry season. If darker coats impose higher thermoregulatory costs, their persistence in subtropical environments like HK raises questions. Darker coats correlate with higher androgens levels and reproductive success (Loehr *et al*., 2008; Bro-Jørgensen & Dabelsteen, 2008; Coulson *et al*., 2011) as well as enhanced heat dissipation through skin evaporation in Holstein cows (Maia *et al*., 2005; Santos *et al*., 2017). Variations in pigmentation relate to behavioural differences (Tucker *et al*., 2008; Stuart-Fox *et al*., 2017), with darker mammals often exhibiting greater dominance, fertility, aggression, and sexual activity (Côté & Festa-Bianchet, 2001; Ducrest *et al*., 2008; Mckinnon & Pierotti, 2010; Roulin, 2014). In giraffes (*Giraffa camelopardalis*), darker males were more likely to live in smaller social groups and to spend more time alone, reflecting different sexual strategies between paler and darker males, with coat colour playing a role as a signal of competitive ability (Castles *et al*., 2019). Similarly, in eland (*Tragelaphus oryx*), acquiring a higher social status led to darkening of facial colouration, highlighting a social function of phenotypic plasticity in coat colour in ungulates (Bro-Jørgensen & Beeston, 2015). However, contrasting evidence exists with paler male Himalayan tahr (*Hemitragus jemlahicus*) being larger, higher ranked and having a higher reproductive success than their darker conspecifics (Lovari *et al*., 2009, 2015; Fattorini *et al*., 2024). Though cattle can discriminate long wavelengths from medium wavelengths (Phillips & Lomas, 2001) suggesting they would be able to discriminate red-based coat colours, further evidence on their ability to discriminate conspecifics based on coat colour and to adjust their behaviour is lacking, but may explain why darker coats persist in this population.

While the coat colour of HK cattle is plastic and exhibits seasonal patterns, most individuals show limited variation in colouration. Cattle with extreme coat colours are more likely to moult, which occurs when the cost of producing a different coat is lower than maintaining a suboptimal colouration (Beltran *et al*., 2018). Consequently, cattle with extreme coats may face higher costs when their pelage becomes maladapted, increasing the likelihood of moulting. Similarly, in springbok, extreme coat colours (black and white) were rare, likely due to variations in thermoregulation costs (Hetem *et al*., 2009). Extreme colours can yield both significant benefits and higher costs, leading most individuals to exhibit average traits that balance these factors. While paler coats can reduce heat stress, darker individuals often enjoy reproductive and immune advantages (Ducrest *et al*., 2008; Coulson *et al*., 2011; Delhey, 2019). Our study addresses a simplified version of Gloger’s rule, examining pigmentation as a whole; however, melanin pigments (*i.e.* eumelanin and phaeomelanin) are known to respond differently to climatic conditions (Delhey, 2019; Marcondes *et al*., 2020). Further investigation into melanism in feral cattle could enhance our understanding of how climate and thermoregulation costs influence the prevalence of coat colours.

In artiodactyls, males tend to be both larger and longer than females (Cassini, 2020; Tombak *et al*., 2024), similar to our findings that HK feral bulls were larger in all body size measures than HK feral cows. Domestication in bovids has led to greater sexual dimorphism with larger males (Polák & Frynta, 2009; McPherson & Chenoweth, 2012). Among cattle, milk breeds are the most sexually dimorphic, while draught breeds are the least dimorphic (Polák & Frynta, 2010). Additionally, cattle breeds from the tropics and subtropics are less sexually dimorphic than breeds from temperate zones (Polák & Frynta, 2010), with Sexual Dimorphism Index (SDI) in South Asian breeds averaging 1.2. Hence, the moderate sexual dimorphism of HK feral cattle (1.07) may result from their South Asian origin and their genetic similarity with other South Asian cattle, along with their history as draught animals. More specifically, SDI in wild bovids averages 1.1 in indicine breeds and Gayal (*Bos frontalis*; Polák & Frynta, 2010). Thus, HK feral cattle not only share genetic relatedness to Asian wild bovids (Barbato *et al*., 2020), we now show they also share phenotypic similarities through moderate sexual dimorphism.

Male bovids usually have longer horns than their female counterparts (Emlen, 2008; Stankowich & Caro, 2009; McPherson & Chenoweth, 2012), as we found in the HK cattle. Sexual selection has been shown to condition horns in males and females differently (Bro-Jørgensen, 2007; Stankowich & Caro, 2009). Horns are primarily used as weapons in male-male competition (Lundrigan, 1996; Berglund *et al*., 1996; Emlen, 2008; McCullough *et al*., 2016), resulting in secondary involvement in mate selection, with females favouring males with longer horns in several bovid species (Berglund *et al*., 1996; Knierim *et al*., 2015). Hence, males may benefit from investing resources into growing long horns to improve their reproductive success, while females may favour other functions such as defence against predators, as found in other bovid species (Bro-Jørgensen, 2007). Female HK feral cattle are more likely to have broken horns than males, which may result from differences in behaviour or susceptibility to environmental factors between males and females but is currently unknown.

## CONCLUSION

The Hong Kong (HK) feral cattle are free roaming in a subtropical environment. We provide the first assessment of the sexual dimorphism in horn and body size and seasonal changes in coat colour of the HK feral cattle. The sexual dimorphism in HK feral cattle is male-biased, and moderate, comparable to other South Asian and draught breeds. Surprisingly, we found that female HK cattle were more likely than males to have broken horns, but literature about the mechanisms underlying these differences are lacking. Extreme coat colours are less common and more susceptible to seasonal changes, with paler individuals in the wet season and darker ones in the dry season, suggesting plastic phenotypes. The main driver of seasonal coat colour appears to be thermoregulation. As cattle are sensitive to heat stress, understanding the mechanism through which mismatched phenotypes remain (*e.g.* darker individuals with higher thermoregulation costs in the wet season) has consequences for management of ruminants in changing environments. Additionally, with the global shift in climate, understanding the drivers of phenotypic plasticity in subtropical cattle improves our knowledge of adaptation in tropical and subtropical free-ranging populations in increasingly warm and unpredictable habitats.

## Acknowledgements

The authors are grateful to the “Sai Kung Bovid Watch”, “Hong Kong Bovid Conservation Association” and the AFCD cattle management team that provided valuable information on the cattle. We are thankful to Yeuk Wan Sharon Lam, Wing Sum Wong, Wang Hin Wong and Claire Giraudet for their valuable contribution to fieldwork and data collection. We are also grateful to Yifu Wang for providing the files for the maps of Hong Kong. We also thank Dieter Lukas and André Vieira Rodrigues for their helpful comments on previous versions of this manuscript.

## SUPPORTING INFORMATION

**Sexual dimorphism and phenotypic plasticity in a feral bovid *Bos taurus* (Artiodactyla: Bovidae).**

*Includes:*

### Tables

Table S1: Description of the eight cattle coat colour scores removed from further analysis due to mode (most frequent value between fore, middle and hind body coat colour) not being obtained.

Table S2: Climate variables extracted from the Hong Kong Observatory (HKO) for each month when coat colour scoring was conducted.

Table S3: Number of female and male cattle scored in each herd and overall number of individuals where at least one phenotype trait could be scored for each herd and country park.

Table S4: Relationships between the five measures of cattle body size (wither height, hip height, body length, hip width and chest depth) used to measure Hong Kong feral cattle body size, as evaluated by Spearman’s correlations.

Table S5: Differences of coat colours (scored among 5 categories in a colour chart) between male and female cattle.

Table S6: Coat colour predicted in relation to temperature (in °C) in Hong Kong feral cattle (n = 253) based on Predicted Probabilities (PP) of Ordinal Logistic Regression.

Table S7: Coat colour predicted in relation to Body Condition Score (BCS) in Hong Kong feral cattle (n = 253) based on Predicted Probabilities (PP) of Ordinal Logistic Regression.

Table S8: Coat colour change magnitude between the three seasons in Hong Kong feral cattle (n = 253) based on Tukey Post-hoc test.

### Figures

Figure S1: Calves (A) were defined as both individuals without horns and individuals of similar appearance to same-aged cows. Juveniles (B) have horn tips and distinct coat patterns.

