## Supplementary material for "Seasonal changes in coat colour and sexual size dimorphism in a subtropical ungulate": Table S1; Table S2; Table S3; Table S4; Table S5; Table S6; Table S7; Table S8; Figure S1

**SUPPORTING INFORMATION**

**Sexual dimorphism and phenotypic plasticity in a feral bovid *Bos taurus* (Artiodactyla: Bovidae).**

*Includes:*

**Tables:**

Table S1: Description of the eight cattle coat colour scores removed from further analysis due to mode (most frequent value between fore, middle and hind body coat colour) not being obtained.

Table S2: Climate variables extracted from the Hong Kong Observatory (HKO) for each month when coat colour scoring was conducted.

Table S3: Number of female and male cattle scored in each herd and overall number of individuals where at least one phenotype trait could be scored for each herd and country park.

Table S4: Relationships between the five measures of cattle body size (wither height, hip height, body length, hip width and chest depth) used to measure Hong Kong feral cattle body size, as evaluated by Spearman’s correlations.

Table S5: Differences of coat colours (scored among 5 categories in a colour chart) between male and female cattle.

Table S6: Coat colour predicted in relation to temperature (in °C) in Hong Kong feral cattle (n = 253) based on Predicted Probabilities (PP) of Ordinal Logistic Regression.

Table S7: Coat colour predicted in relation to Body Condition Score (BCS) in Hong Kong feral cattle (n = 253) based on Predicted Probabilities (PP) of Ordinal Logistic Regression.

Table S8: Coat colour change magnitude between the three seasons in Hong Kong feral cattle (n = 253) based on Tukey Post-hoc test.

**Figures:**

Figure S1: Calves (A) were defined as both individuals without horns and individuals of similar appearance to same-aged cows. Juveniles (B) have horn tips and distinct coat patterns.

**Table S1: Description of the eight cattle coat colour scores removed from further analysis due to mode (most frequent value between fore, middle and hind body coat colour) not being obtained.**

| **Cow ID** | **Sex** | **Group** | **Month** | **Season** | **Year** | **Fore Coat Colour** | **Middle Coat Colour** | **Hind Coat Colour** |
| --- | --- | --- | --- | --- | --- | --- | --- | --- |
| Apple | F | Kuk Po Lo Wai | August | Wet | 2022 | Red | Pale | Grey |
| 97 | F | Chong Hing | August | Wet | 2022 | Dark | Red | Grey |
| Mike | M | Ham Tin | August | Wet | 2022 | Dark | Red | Grey |
| Bob | M | Ham Tin | August | Wet | 2022 | Dark | Red | Grey |
| 726 | F | Pak Lap | February | Dry | 2023 | Grey | Dark | Red |
| Elf | F | Kuk Po Lo Wai | February | Dry | 2023 | Dark | Red | Grey |
| 718 | M | Chong Hing | August | Wet | 2023 | Grey | Red | Pale |
| Wanda | F | Tap Mun | January | Dry | 2024 | Dark | Red | Pale |

**Table S2: Climate variables extracted from the Hong Kong Observatory (HKO) for each month when coat colour scoring was conducted.**

| **Season** | **Month** | **Mean daily temperature (C°)** | **Relative humidity (%RH)** | **Total rainfall (mm)** | **Total bright sunshine duration (h)** | **Mean daily global solar radiation (MJ per m²)** |
| --- | --- | --- | --- | --- | --- | --- |
| Wet | August 2022 | 28.8 | 82 | 614.8 | 167.7 | 16.22 |
| Dry | January 2023 | 17.0 | 67 | 18.2 | 134.1 | 11.44 |
|  | February 2023 | 18.9 | 73 | 1.6 | 163.8 | 14.64 |
| Intermediate | March 2023 | 21.3 | 76 | 70.3 | 156.8 | 13.13 |
|  | April 2023 | 23.6 | 82 | 77.5 | 92.3 | 11.02 |
| Wet | May 2023 | 26.6 | 81 | 182.8 | 131.9 | 14.59 |
|  | June 2023 | 29.2 | 83 | 490.9 | 147.4 | 15.17 |
|  | July 2023 | 30.1 | 78 | 175.2 | 219.2 | 19.13 |
|  | August 2023 | 29.7 | 79 | 140.7 | 166.4 | 16.02 |
|  | September 2023 | 28.5 | 81 | 1067.1^#^ | 170.5 | 15.31 |
|  | October 2023 | 26.4 | 76 | 546.0 | 138.9 | 12.25 |
| Intermediate | November 2023 | 23.5 | 69 | 3.3 | 208.2 | 14.57 |
| Dry | December 2023 | 19.1 | 70 | 0.9 | 136.0 | 10.70 |
|  | January 2024 | 17.9 | 72 | 6.7 | 169.8 | 12.42 |

*# occurrence of an extreme flooding following a rainstorm lasting 16 hours on 7^th^ to 8^th^ of September 2023 and exceeding normal rainfall in September by 3 times.*

**Table S3:** **Number of female and male cattle scored in each herd and overall number of individuals where at least one phenotype trait could be scored for each herd and country park.**

| **Country Park** | **Herd** | **Sex** | **Number of individuals scored** | **Total Per herd** | **Total per country park** |
| --- | --- | --- | --- | --- | --- |
| Crooked Island | Kat O | F | 4 | 5 | 5 |
|  |  | M | 1 |  |  |
| Grass Island | Tap Mun | F | 10 | 15 | 15 |
|  |  | M | 5 |  |  |
| Lantau North | Lin Yan Monastery | F | 8 | 13 | 33 |
|  |  | M | 5 |  |  |
|  | Ngong Ping | F | 1 | 3 |  |
|  |  | M | 2 |  |  |
|  | San Shek Wan | F | 9 | 15 |  |
|  |  | M | 6 |  |  |
|  | Tai O | F | 1 | 2 |  |
|  |  | M | 1 |  |  |
| Lantau South | Mui Wo | F | 4 | 19 | 34 |
|  |  | M | 15 |  |  |
|  | Shek Pik | F | 4 | 7 |  |
|  |  | M | 3 |  |  |
|  | Tong Fuk | F | 4 | 8 |  |
|  |  | M | 4 |  |  |
| Ma On Shan | Ngong Ping Ma On Shan | F | 4 | 7 | 17 |
|  |  | M | 3 |  |  |
|  | Sai Kung Town | F | 0 | 5 |  |
|  |  | M | 5 |  |  |
|  | Yan Yee Road | F | 5 | 5 |  |
|  |  | M | 0 |  |  |
| Plover Cove | Kai Kuk Shue Ha | F | 0 | 2 | 27 |
|  |  | M | 2 |  |  |
|  | Kuk Po Lo Wai | F | 12 | 20 |  |
|  |  | M | 8 |  |  |
|  | So Lo Pun | F | 1 | 4 |  |
|  |  | M | 3 |  |  |
|  | YungShueAu | F | 0 | 1 |  |
|  |  | M | 1 |  |  |
| Sai Kung East | Chong Hing | F | 26 | 43 | 117 |
|  |  | M | 17 |  |  |
|  | Ham Tin | F | 7 | 18 |  |
|  |  | M | 11 |  |  |
|  | Long Ke | F | 3 | 9 |  |
|  |  | M | 6 |  |  |
|  | Pak Lap | F | 16 | 29 |  |
|  |  | M | 13 |  |  |
|  | Sai Wan | F | 4 | 10 |  |
|  |  | M | 6 |  |  |
|  | Tung Wan | F | 0 | 1 |  |
|  |  | M | 1 |  |  |
|  | Wong Shek | F | 5 | 7 |  |
|  |  | M | 2 |  |  |
| Sai Kung West | Cheung Sheung | F | 5 | 6 | 16 |
|  |  | M | 1 |  |  |
|  | Hoi Ha | F | 1 | 2 |  |
|  |  | M | 1 |  |  |
|  | Lai Chi Chong | F | 5 | 8 |  |
|  |  | M | 3 |  |  |
| Sheung Shui | Sheung Shui | F | 12 | 16 | 16 |
|  |  | M | 4 |  |  |
| Shing Mun | Grassy Hill | F | 10 | 20 | 27 |
|  |  | M | 10 |  |  |
|  | Shing Mun | F | 4 | 7 |  |
|  |  | M | 3 |  |  |
| Tai Lam | Tsin Fai Tong | F | 6 | 10 | 10 |
|  |  | M | 4 |  |  |

**Table S4: Relationships between the five measures of cattle body size (wither height, hip height, body length, hip width and chest depth) used to measure Hong Kong feral cattle body size, as evaluated by Spearman’s correlations.**

|  |  |  | **n** | **Spearman's rho** | **p-value** | **Lower 95%CI** | **Upper 95%CI** |
| --- | --- | --- | --- | --- | --- | --- | --- |
| Wither Height | - | Hip Height | 228 | 0.866 | < .001 | 0.813 | 0.905 |
| Wither Height | - | Body Length | 228 | 0.830 | < .001 | 0.762 | 0.882 |
| Wither Height | - | Hip Width | 231 | 0.542 | < .001 | 0.443 | 0.631 |
| Wither Height | - | Chest Depth | 234 | 0.920 | < .001 | 0.884 | 0.944 |
| Hip Height | - | Body Length | 227 | 0.810 | < .001 | 0.732 | 0.865 |
| Hip Height | - | Hip Width | 232 | 0.656 | < .001 | 0.560 | 0.734 |
| Hip Height | - | Chest Depth | 232 | 0.824 | < .001 | 0.757 | 0.876 |
| Body Length | - | Hip Width | 233 | 0.643 | < .001 | 0.546 | 0.724 |
| Body Length | - | Chest Depth | 234 | 0.824 | < .001 | 0.753 | 0.878 |
| Hip Width | - | Chest Depth | 238 | 0.546 | < .001 | 0.439 | 0.633 |

*Body lengths significantly differed from normality as assessed by Shapiro-Wilk test.* *Confidence intervals are based on 1000 bootstrap replicates. N indicates the number of scores compared in each correlation.*

**Table S5: Differences of coat colours (scored among 5 categories in a colour chart) between male and female cattle. Grey background indicates months where males and female coat colour distribution was similar, based on Χ² test.**

| **Month** | **Χ²** | **N** | **p-value** |
| --- | --- | --- | --- |
| January | 28.998 | 349 | <0.0001 |
| February | 19.639 | 169 | <0.0001 |
| March | 22.485 | 155 | <0.0001 |
| April | 12.535 | 162 | 0.014 |
| May | 8.631 | 185 | 0.071 |
| June | 2.964 | 169 | 0.564 |
| July | 11.898 | 161 | 0.018 |
| August | 28.960 | 338 | <0.0001 |
| September | 6.462 | 164 | 0.167 |
| October | 8.921 | 167 | 0.063 |
| November | 19.011 | 170 | <0.0001 |
| December | 11.170 | 168 | 0.025 |
| Total | 156.207 | 2357 | <0.0001 |

*N indicates the number of cattle scored per month*

**Table S6: Coat colour predicted in relation to temperature (in °C) in Hong Kong feral cattle (n = 253) based on Predicted Probabilities (PP) of Ordinal Logistic Regression.**

| **Temperature (°C)** | **Pale** | **Red** | **Grey** | **Dark** | **Black** |
| --- | --- | --- | --- | --- | --- |
| **15** | 11.57 | 31.04 | 7.1 | 19.76 | 30.5 |
| **20** | 15.36 | 35.38 | 7.09 | 18.11 | 24.03 |
| **25** | 20.12 | 38.71 | 6.72 | 15.86 | 18.57 |
| **30** | 25.89 | 40.58 | 6.05 | 13.34 | 14.12 |
| **35** | 32.64 | 40.68 | 5.21 | 10.84 | 10.59 |
| **Mean** | 21.11 | 37.27 | 6.43 | 15.58 | 19.56 |

*PP were converted to %, with each column summing up to 100; Temperature, though a numerical variable in the model, is presented for the temperature range of Hong Kong.*

**Table S7: Coat colour predicted in relation to Body Condition Score (BCS) in Hong Kong feral cattle (n = 253) based on Predicted Probabilities (PP) of Ordinal Logistic Regression.**

| **BCS** | **Pale** | **Red** | **Grey** | **Dark** | **Black** |
| --- | --- | --- | --- | --- | --- |
| **3** | 16.39 | 35.25 | 6.99 | 17.66 | 23.68 |
| **4** | 19.01 | 37.11 | 6.81 | 16.47 | 20.59 |
| **5** | 21.93 | 38.55 | 6.53 | 15.16 | 17.80 |
| **6** | 25.17 | 39.52 | 6.17 | 13.80 | 15.32 |
| **7** | 28.71 | 39.98 | 5.74 | 12.42 | 13.12 |
| **Mean** | 22.24 | 38.08 | 6.44 | 15.10 | 18.10 |

*PP were converted to %, with each column summing up to 100. While BCS were scored on a 9-point scale, the range selected here represented the distribution of BCS found in HK feral cattle.*

**Table S8: Coat colour change magnitude between the three seasons in Hong Kong feral cattle (n = 253) based on Tukey Post-hoc test. Bold font indicates significant comparisons.**

|  | | | **Mean Difference** | **95% Confidence Interval** | | | **SE** | **t** | **p-value** |
| --- | --- | --- | --- | --- | --- | --- | --- | --- | --- |
| **Wet** | **-** | **Intermediate** | **0.200** | **0.048** | **-** | **0.352** | **0.065** | **3.094** | **0.006** |
| Wet | - | Dry | 0.146 | -0.011 | - | 0.303 | 0.067 | 2.176 | 0.08 |
| Intermediate | - | Dry | -0.054 | -0.185 | - | 0.076 | 0.056 | -0.976 | 0.98 |

*p-values were adjusted using Bonferroni correction for multiple comparisons (here 3).*

|  | 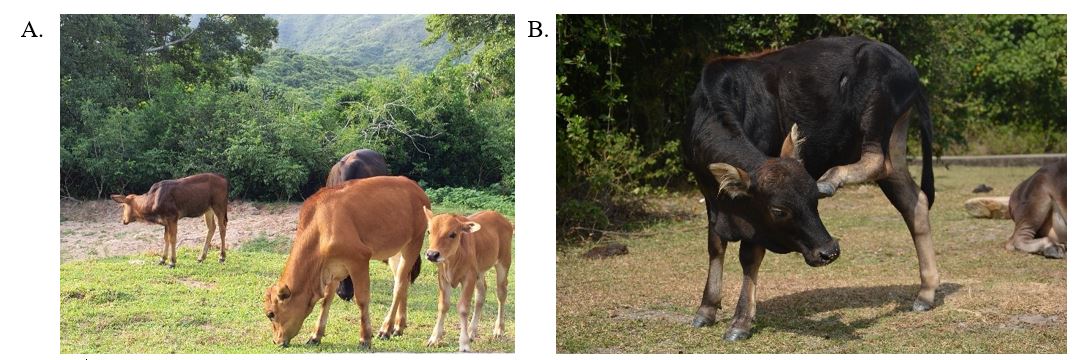 |
| --- | --- |

**Figure S1: Calves (A) were defined as both individuals without horns and individuals of similar appearance to same-aged cows. Juveniles (B) have horn tips and distinct coat patterns.**

*Photo credit: Nicolas Brualla (left), George Hodgson (right)*
